# PARP14 is a PARP with both ADP-ribosyl transferase and hydrolase activities

**DOI:** 10.1101/2023.06.25.546318

**Authors:** Nina Đukić, Øyvind Strømland, Deeksha Munnur, Kang Zhu, Marion Schuller, Chatrin Chatrin, Pulak Kar, Johannes Gregor Matthias Rack, Domagoj Baretić, Herwig Schüler, Sven Wijngaarden, Dmitri V. Filippov, Sumana Sanyal, Rebecca Smith, Dragana Ahel, Ivan Ahel

## Abstract

PARP14 is a mono-ADP-ribosyl transferase involved in the control of immunity, transcription and DNA replication stress management. However, little is known about the ADP-ribosylation activity of PARP14, including its substrate specificity or how PARP14-dependent ADP-ribosylation is reversed. Here we show that PARP14 is dual function enzyme with both ADP-ribosyl transferase and hydrolase activity acting on both protein and nucleic acid substrates. In particular, we show that the PARP14 macrodomain 1 is an active ADP-ribosyl hydrolase. We also demonstrate hydrolytic activity for the first macrodomain of PARP9. We reveal that expression of a PARP14 mutant with the inactivated macrodomain 1 results in a dramatic increase in mono(ADP-ribosyl)ation of proteins in human cells, including PARP14 itself and antiviral PARP13. Moreover, we demonstrate that the closely related hydrolytically active macrodomain of SARS2 Nsp3, Mac1, efficiently reverses PARP14 ADP-ribosylation *in vitro* and in cells, supporting the evolution of viral macrodomains to counteract PARP14-mediated antiviral response.

**Teaser:** PARP14 is an antiviral PARP that combines ADP-ribosylation writer, reader and eraser functions in one polypeptide.

## Introduction

Cells must quickly adapt to both internal and external changes. This could be due to internal pressures such as changes in metabolic demands that require alteration in transcription or protein translation, DNA replication or DNA damage repair, as well as from external pressures such as invasion of pathogenic bacteria and viruses. In order for such changes to occur efficiently, cells have developed a number of signaling pathways to transduce signals rapidly. This often involves post-transcriptional or post-translational modification of nucleic acids and proteins respectively. One such signaling type is the ADP-ribosylation (ADPr). ADPr has been shown to target both proteins (*1*) and nucleic acids, including RNA and DNA (*2*), with modification on proteins able to occur on different amino acid acceptors including serine, glutamate and arginine (*3-5*).

Like in most signaling pathways, there are proteins required for the writing, reading and reversal of ADPr. The largest known family of proteins that are responsible for the addition of ADP-ribose to proteins or nucleic acids are the PARPs. PARPs function by transferring ADP-ribose from the dinucleotide NAD^+^ onto a target, releasing nicotinamide. These ADP-ribose moieties can either be added as a single moiety, known as mono-ADP-ribosylation, or it can be attached as long branched polymers, so called poly(ADP-ribosyl)ation (*6*). A number of different reader domains have been shown to recognise and bind ADP-ribose including macrodomains which primarily recognise either mono-ADP-ribose or the terminal ADP-ribose moiety of a polymer, or WWE domains and, PAR-binding zinc finger (PBZ) that specifically recognise poly(ADP-ribose) (*7-10*). Finally, there are eraser proteins involved in the removal of ADP-ribose, primarily ADP-ribosylhydrolases (ARHs), or hydrolytic macrodomains, of which four, poly(ADP-ribosyl) glycohydrolase (PARG), MacroD1, MacroD2 and TARG1, have been described in humans (*11*).

While there have been strides taken in understanding how several members of the PARP family function, there are still outstanding questions regarding the specificity of their ADP-ribosylation activity, as well as the hydrolases that remove their modification. The best understood aspect of ADPr signaling in humans is its role in the DNA damage response. Here PARP1 or PARP2 will recognise and bind to DNA breaks (*12*) where they interact with their auxiliary factor HPF1 (*13*). Together they ADP-ribosylate proteins around the break site, including PARP1 itself, on serine residues (*14, 15*). The modified proteins can be also then recognised by ADP-ribose-binding repair proteins such as ALC1, XRCC1 and APLF which are required for efficient DNA repair (*16*). Following the initial ADP-ribose signaling at DNA breaks, the ADP-ribose moieties are efficiently removed by PARG, which degrades long polymers of ADP-ribose, while the final ADP-ribose on serine residues is removed by ARH3 (*17*). Recently, we have also begun to understand the complexity of ADPr signaling on nucleic acids in bacterial systems, where a PARP-like protein, DarT, catalyses the ADP-ribosylation of thymidine bases which can be efficiently reversed by the hydrolytic macrodomain DarG (*18, 19*). Several human PARPs have been suggested to ADP-ribosylate 5’and 3’phosphorylated single-stranded DNA and RNA (*20-22*) (*23*). This is reversed by endogenous ADP-ribosyl hydrolases, including PARG, TARG1, MACROD1, MACROD2 and ARH3 (*20, 21*).

Despite the wealth of knowledge on ADPr and PARP1, much less is known about most of the other human PARPs. One poorly understood subgroup are the interferon-induced “antiviral PARPs” which include PARP7, 9, 10, 11, 12, 13, 14 and 15. Several members of this class have been reported to modify both protein and nucleic acid substrates (*20-22*), while PARP9 and PARP13 are reported to be catalytically inactive (*3*). These antiviral PARPs are interferon-inducible and were shown to confer resistance to a range of viruses, including coronaviruses, influenza, HIV and Ebola (*24-27*). These PARPs are also under positive evolutionary selection, strongly suggesting co-evolution with viruses as a consequence of host-virus conflicts (*28*). The largest of all the human PARPs is PARP14. The N-terminus of PARP14 harbours three RRM domains as well as 8 putative KH domains which may bind of RNA or DNA, in addition to three ADPr-binding macrodomains (Fig. 1A). The C-terminus of PARP14 contains a WWE domain, followed by the catalytic ADP-ribosyl transferase (ART) domain.

**Figure 1:**
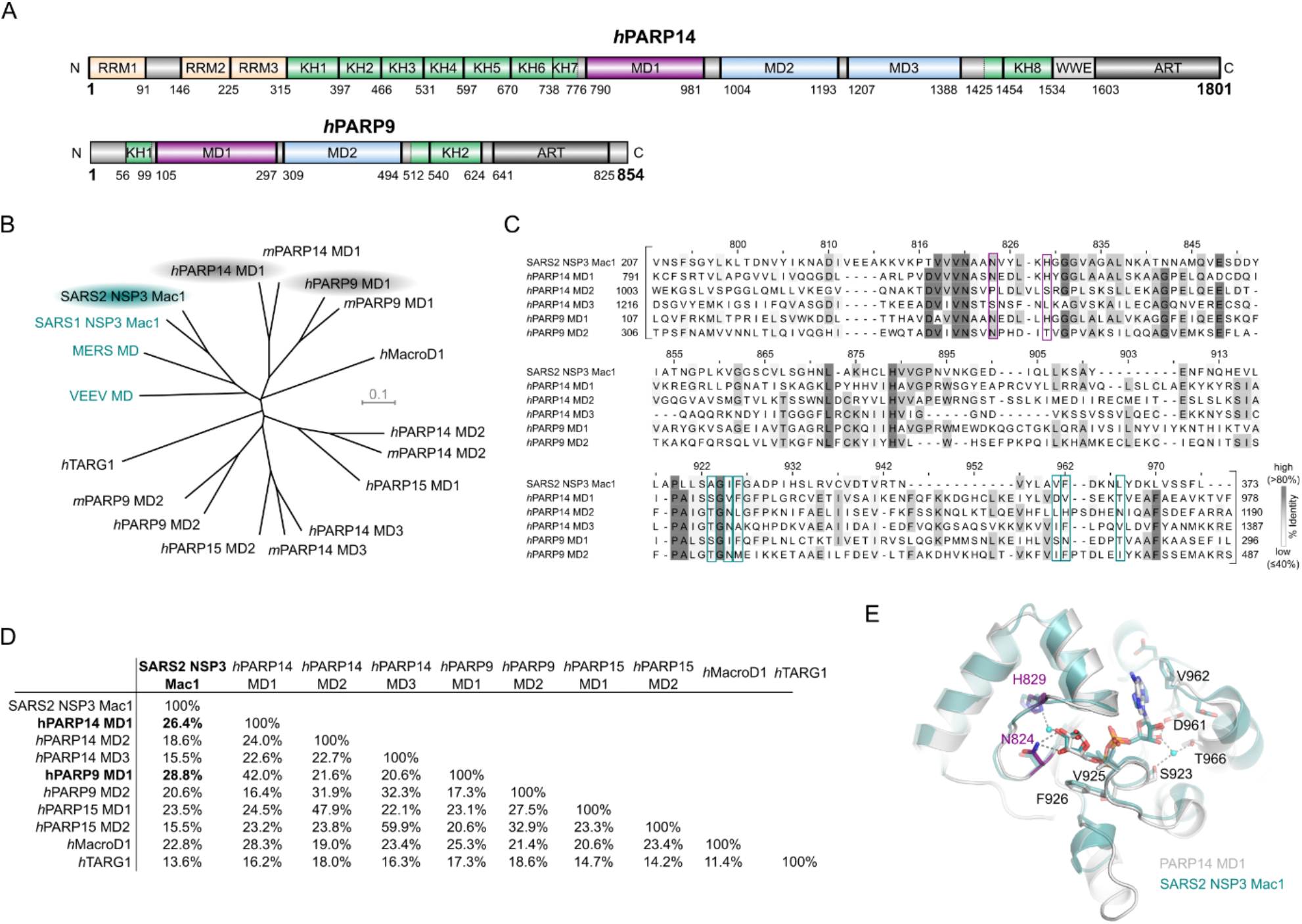
PARP14 and PARP9 macrodomain 1 are similar to SARS2 Mac1. **(A)** Domain architecture of human PARP14 and PARP9. **(B)** Unrooted phylogenetic tree of human and mouse macrodomains including of PARP9, 14 and 15, and viral macrodomains (highlighted in cyan). **(C)** Multiple sequence alignment showing conservation of catalytic residues (magenta-framed) and residues involved in ADP-ribose coordination (cyan-framed) of human PARP14 and PARP9 macrodomains in comparison to SARS2 Nsp3 Mac1. Numbers on top of the residues refer to human PARP14 MD1. **(D)** Pairwise sequence identity comparison of SARS2 Nsp3 Mac1 and human macrodomains. **(E)** Crystal structure overlay of PARP14 MD1 (PDB ID: 3Q6Z) and SARS2 Nsp3 Mac1 (PDB ID: 7KQP) both in complex with ADP-ribose (r.m.s.d of 1.02Å over 214 C^α^). The catalytic residues are highlighted in magenta.

PARP14 has been reported to regulate several different pathways involved in immunity, inflammation and genome stability. PARP14 was initially shown to be involved in transcriptional regulation, acting as a molecular switch of IL-4-regulated genes (*29*). In basal conditions, PARP14 represses gene transcription by binding to IL-4 responsive promotor and recruiting histone deacetylase-2/3 (HDAC2/3). In contrast, under IL-4 stimulated conditions, PARP14 is activated leading to the dissociation of HDAC2/3 from the promoter regions. Consequently, this allows the binding of the transcription factor STAT6, and other transcription co-factors, to their target genes and allows efficient gene transcription (*30*). PARP14 has also been shown to regulate transcription in response to IFNγ stimulation. Specifically, PARP14 has been suggested to ADP-ribosylate STAT1, inhibiting phosphorylation of STAT1 and subsequent activation of pro-inflammatory gene expression. With its role in regulating genomic stability, PARP14 was identified a component of homologous recombination (HR) DNA repair machinery (*31, 32*). Recently, PARP14 has been reported to regulate the replication stress response through recruiting Mre11 nuclease to stalled replication forks (*33*).

PARP14 has also been reported to play an important role in the antiviral response (*26, 34, 35*). In the context of coronavirus infection, PARP14 is required to enhance type I interferon production and restrict replication of murine hepatitis virus (MHV), a model coronavirus (*26*). To combat the antiviral activity of these PARPs, severe acute respiratory syndrome coronavirus (SARS-CoV) contains a hydrolytic macrodomain (*36, 37*) within the Non-structural Protein 3 (Nsp3) which has been suggested to remove PARP14 mediated ADPr (*38*). Importantly, Nsp3 macrodomain 1 (Mac1) is critical for virus replication *in vivo* and viruses with mutated or absent macrodomains are unable to hydrolyse host ADP-ribosylation and therefore associated with reduced viral loads and increased sensitivity to IFN-I treatment in PARP14-proficient cells (*26, 27*). However, the exact mechanism of activation and the molecular substrates of PARP14 and the viral macrodomains are still poorly characterised. Notable, PAPR14 has been shown to efficiently modify itself both in its catalytic region and isolated MD2 and MD3, but not MD1 (*3, 38*) and ADP-ribosylation of endogenous PARP14 on acidic residues has also been detected in IFNγ-stimulated primary human macrophages (*39*).

PARP14s closest relative, PARP9, is also induced by interferon stimulation and is expressed from the same genetic locus as PARP14 (Chromosome 3q21.1). Structurally, PARP9 is similar to PARP14 with two KH domains and two macrodomains at the N-terminus and a C-terminal ART domain (Fig. 1A). No ADP-ribosyl transferase activity of PARP9 has been detected to date, presumably due to the lack of several essential catalytic amino acids within the ART domain (*3*).

Previous studies have suggested that the Mac1 of SARS-CoV-2 (SARS2) Nsp3 is closely related to macrodomain 1 of PARP9 and PARP14 (*38*). Given these similarities, we sought to examine if PARP9 and PARP14 MD1 share the same hydrolytic function. Here we show that these macrodomains are indeed active hydrolases and can remove ADPr from both protein and nucleic acid substrates. We also show that PARP14 modifies proteins in cells by mono-ADPr and that this ADPr is reversed by its own macrodomain. Thus, we show that PARP14 is a PARP that acts both as a transferase, and as a hydrolase. We further show that SARS2 Mac1 can reverse PARP14 dependent ADPr.

## Results

### PARP14 and PARP9 macrodomain 1 exhibit ADP-ribosyl hydrolase activity on protein substrates

PARP14, the largest of the human PARPs belonging to the macrodomain-containing PARPs, together with PARP9 and PARP15, is a potent mono (ADP-ribosyl) transferase (*3, 38, 40*) (Fig. 1A). PARP14 ADP-ribosylation activity is efficiently reversed *in vitro* by SARS2 Nsp3 Mac1 (*38*), however, human endogenous hydrolases that can reverse PARP14 modification remain elusive. To identify potential human hydrolases with activity towards PARP14 mediated ADP-ribosylation, we compared human macrodomains to Mac1. Phylogenetic analysis suggests that while there is an obvious homology to the known hydrolases such as MacroD1, the closest orthologs are the first macrodomain of PARP14 (PARP14 MD1) and the first macrodomain of PARP9 (PARP9 MD1) (Fig. 1B, C and D) while macrodomains 2 and 3 of PARP14 and macrodomain 2 of PARP9 are more diverged. Comparison of ADP-ribose complex structures of PARP14 MD1 and Mac1 reveals that the residues important for distal ribose coordination are also structurally conserved, suggesting that PARP14 MD1 and Mac1 potentially share same catalytic ability and functional context (Fig. 1D and E).

These observations prompted us to investigate the potential ADP ribosyl hydrolase activities of the macrodomains in PARP9 and PARP14. First, we used the automodified PARP14 catalytic fragment (WWE-CAT) as a model substrate and the isolated macrodomains derived from PARP14 (MD1-3) as described previously (*38*). PARP14 MD1 and SARS2 Mac1 have notable hydrolysis activity on automodified PARP14 WWE-CAT, while MD2 and MD3 did not exhibit activity (Fig. 2A). To strengthen our finding, we introduced a point mutation G832E in PARP14 MD1 which sterically blocks the active site in macrodomain hydrolases for ADP-ribose binding (*41*). As predicted the mutation diminished the catalytic activity of PARP14 MD1 (Fig. 2A). Mutating the corresponding residue in PARP14 MD2 had no discernible effect (Fig. 2A). PARP14 MD3 has also been reported to be robustly modified by PARP14 (Fig. 2A) (*38*). We used this to our advantage and examined if PARP14 MD1 could hydrolyse trans modified PARP14 MD3. Indeed, we were able to observe that PARP14 MD1 and SARS2 Nsp3 Mac1 efficiently reversed PARP14 MD3 ADP-ribosylation (Fig. 2B) while PARP14 MD2 and PARP14 MD3 had no discernible effect on the ADP ribosylation level of trans modified PARP14 MD3.

**Figure 2:**
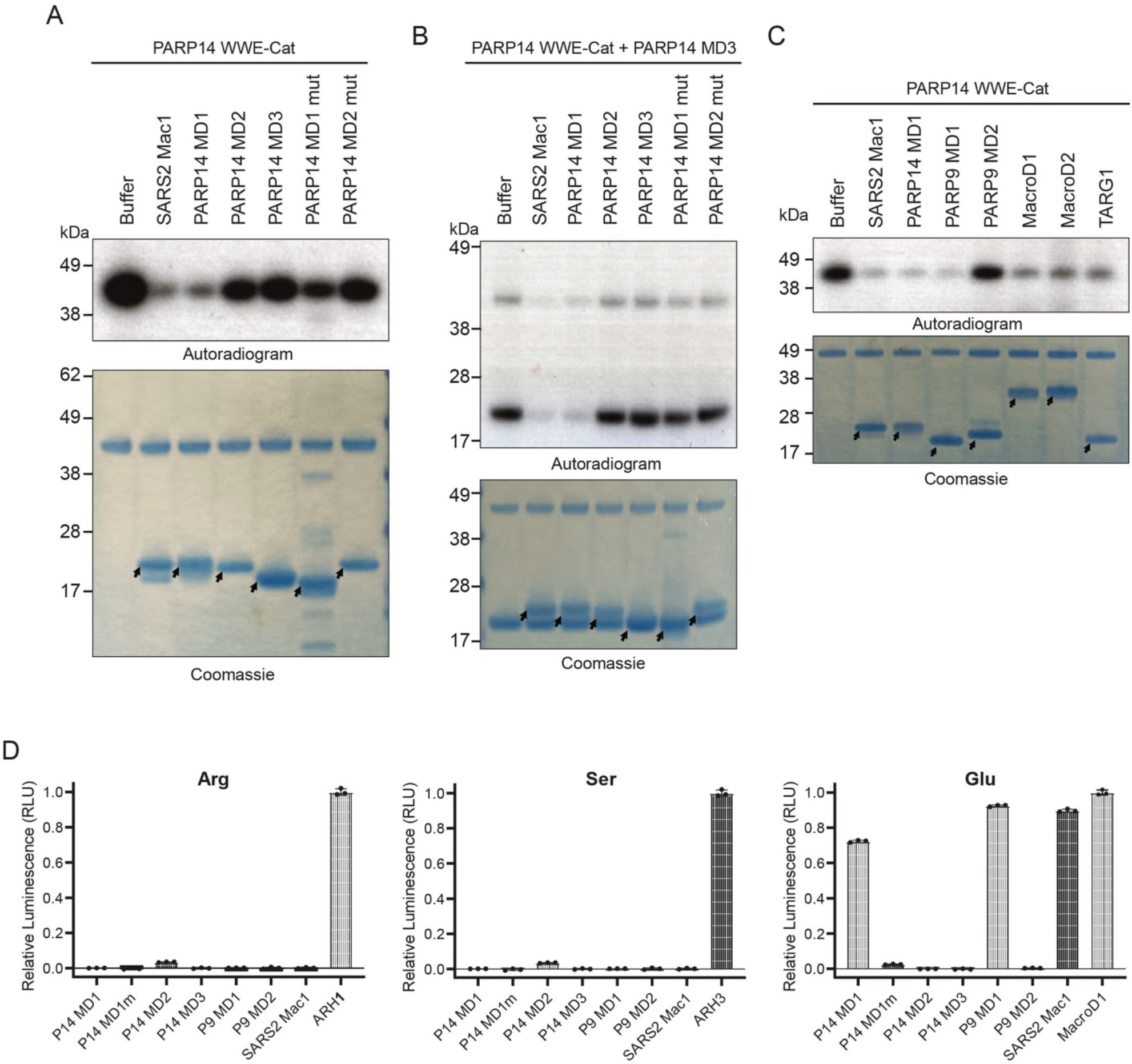
PARP14 and PARP9 MD1 reverse glutamate-linked PARP14 auto- and trans-ADP-ribosylation. **(A)** PARP14 WWE-CAT was auto-ADP-ribosylated using NAD^+^ spiked with ^32^P NAD^+^. The ADP-ribosyl hydrolysis activity of PARP14 MD1, MD2, MD3, MD1mut (G823E), MD2mut (G1044E) and SARS2 Nsp3 Mac1 (SARS2 Mac1) was assessed upon incubation with automodified PARP14 WWE-CAT. **(B)** PARP14 WWE-CAT and PARP14 MD3 were auto- and trans-ADP-ribosylated, respectively, using NAD^+^ spiked with ^32^P NAD^+^. The trans-ADP-ribosylation hydrolysis activity of PARP14 MD1, MD2, MD3, MD1mut, MD2mut and SARS2 Mac1 was assessed upon incubation with the trans modified PARP14 MD3 and auto modified PARP14 WWE-CAT. **(C)** PARP14 WWE-CAT was auto-ADP-ribosylated using NAD^+^ spiked with ^32^P NAD^+^. The ADP-ribosyl hydrolysis activity of SARS2 Mac1, PARP14 MD1, PARP9 MD1, PARP9 MD2, MacroD1, MacroD2 and TARG1 was determined upon incubation with automodified PARP14 WWE-CAT. Samples in (A, B and C) were analysed by Coomassie brilliant blue staining and autoradiography. The arrows show the position of the indicated macrodomains **(D)** Hydrolysis of arginine-, serine- and glutamate linked mono(ADP-ribosyl)ation on synthetic peptides by PARP14 MD1, PARP14 MD1mut, PARP14 MD2, PARP14 MD3, PARP9 MD1, PARP9 MD2 and SARS2 Mac1. Briefly, the released ADP-ribose was converted by NUDT5 to AMP, which subsequently was detected by luminescence using the AMP-Glo™ assay (Promega). Samples are background corrected and normalized to the positive control, ARH1 for arginine, ARH3 for serine and MacroD1 for glutamate. The data represents mean values ± SD measured in triplicates.

PARP14 is involved in macrophage activation whereby gene expression enabling the defense against pathogens is induced. In particular, PARP14 has been suggested to ADP-ribosylate STAT1α, preventing STAT1α phosphorylation which is essential for STAT1α to drive transcription of pro-inflammatory genes (*42*). Conversely, PARP9 has been reported to antagonize the activation by inhibiting the ADP-ribosylation of STAT1α (*42*). Given the similarity between PARP9 MD1, PARP14 MD1 and SARS2 Mac1 and the reported role of PARP9 in antagonizing PARP14 ADP-ribosylation we investigated if PARP9 MD1 can reverse PARP14 automodification. Indeed, PARP9 MD1 could efficiently reverse PARP14 ADP-ribosylation while PARP9 MD2 had no observable effect (Fig. 2C). We also tested several known human hydrolases and saw that MacroD1, MacroD2 and TARG1, known to remove glutamate linked ADP-ribose from target proteins (*19, 43*), exhibited significant hydrolytic activity on automodified PARP14 (Fig. 2C) suggesting that the PARP14 derived ADP-ribosylation could be glutamate linked. To investigate the amino acids specificity of the PARP9 MD1 and PARP14 MD1 we assessed their activity against chemically synthesised defined ADP-ribosylated peptide substrates modified on serine, arginine and glutamate respectively (Fig. 2D). We observed that PARP14 MD1 and PARP9 MD1 are specifically active on Glu-ADPr similar to SARS2 Mac1 and MacroD1. PARP14 MD1 and PARP9 MD1 showed no activity on serine and arginine linked peptides in contrast to the cognate hydrolases ARH3 (*17*) and ARH1 (*44*).

### PARP14 and PARP9 macrodomain 1 exhibit ADP-ribosylhydrolase activity on nucleic acid substrates

The domain architecture of PARP14 suggest that it is tightly linked to nucleic acids since it harbours three RRM domains and eight KH domains which all are putative single stranded RNA or DNA binders (Fig. 1A). Furthermore, PARP14 is interferon induced and plays a role in the innate immune response against viruses (*35*). Interferon induced antiviral PARPs, such as PARP10 and PARP11, ADP-ribosylate 5’and 3’phosphorylated single-stranded (ss) RNA and DNA (*21*). Based on these observations, we postulated that PARP14 MD1 could reverse ssDNA and ssRNA ADP-ribosylation. Single-stranded RNA phosphorylated at either the 5’- or at 3’- were ADP-ribosylated with PARP14 and then used as potential substrates for PARP14 macrodomains (Fig. 3A-C, lane 2). PARP14 MD1 efficiently reversed the ADP-ribosylation from both RNA substrates and, as expected, PARP14 MD2 and MD3 did not (Fig. 3A and B). SARS2 Mac1, as reported previously, could also remove the modification (Fig. 1 A and B) (*21*). Next, we tested if PARP14 MD1 could also reverse the ADP-ribosylation of 5’phosphorylated ssDNA. As for ssRNA, PARP14 MD1 reversed the ADP-ribosylation on ssDNA while PARP14 MD2 and MD3 did not (Fig. 3C). We also determined if PARP9 MD1 can reverse ADP-ribosylation of 5’phosphorylated ssDNA finding that PARP9 MD1 but not MD2 reversed the modification (Fig. 3D). As reported previously MacroD1 could also remove ADP-ribosylation of 5’phosphorylated ssDNA while the unrelated human hydrolase ARH1 could not. Altogether, our data identified two new ADP-ribosylhydrolases in humans (MD1 of PARP9 and PARP14) and demonstrates that PARP14 represents a PARP enzyme that can reverse its own modification.

**Figure 3.**
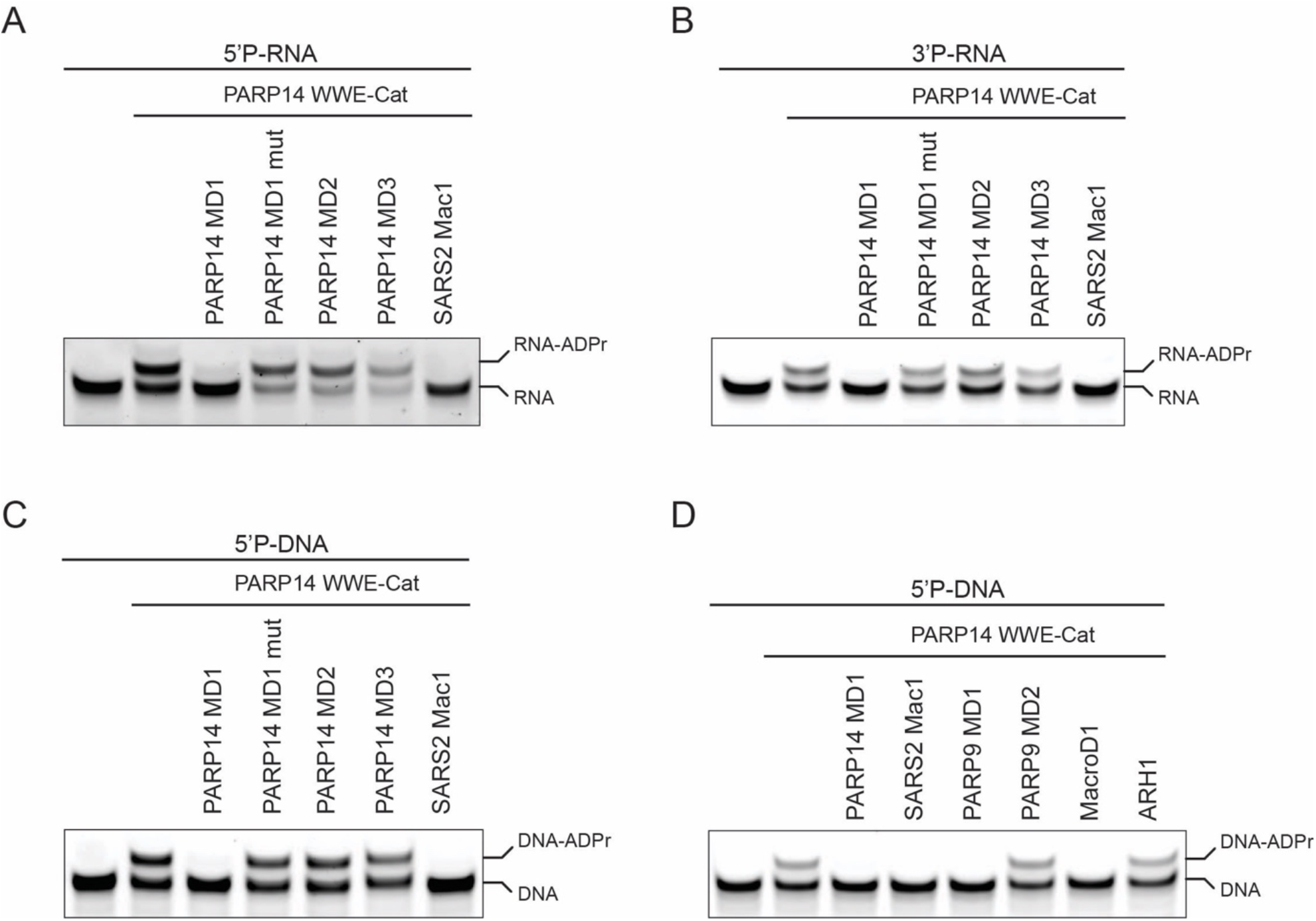
PARP14 and PARP9 MD1 reverse ADP-ribosylation of single stranded RNA and DNA. **(A)** ssRNA with 5’phosphate and 3’Cy3, **(B)** ssRNA with 3’phosphate and 5’Cy3 and **(C)** ssDNA with 5’phosphate and 3’Cy3 were ADP-ribosylated using PARP14 WWE-CAT. Subsequently, the ADP-ribosylation was hydrolysed by treating the modified oligos with PARP14 MD1, MD1mut, MD2, MD3 and SARS CoV2 Mac1. **(D)** ssDNA with 5’phosphate and 3’Cy3 was ADP-ribosylated using PARP14 WWE-CAT. Following, the ADP ribose modification was hydrolysed by subjecting the ADP-ribosylated oligo to PARP14 MD1, SARS CoV2 Mac1, PARP9 MD1 and MD2, MacroD1 and ARH1.

### PARP14 shows ADP-ribosyl transferase and hydrolase activity on different cellular substrates

We next studied PARP14 activity in human cells. To do so, we transiently transfected 293T cells with YFP-tagged full-length PARP14 WT, PARP14 R1699A catalytic mutant, PARP14 G832E MD1 and PARP14 G1044E MD2 mutant, and examined changes in protein mono(ADP-ribosyl)ation in cell extracts using a mono-ADPr specific antibody (Fig. 4A). Overexpression of WT PARP14, but not the R1699A ADP-ribosylation deficient mutant, resulted in a modest increase of mono-ADP-ribosylation, indicating that this mutant is devoid of catalytic activity. On the other hand, we observed a dramatic increase in mono(ADP-ribosyl)ation of a variety of protein sizes when we overexpressed the PARP14 MD1 mutant, suggesting that PARP14 modifies a number of different proteins in the cells. We also noted that a single dominant ADPr signal around 50 kDa was induced specifically in cells overexpressing PARP14 MD2 mutant. In all cases, treatment with a specific PARP14 inhibitor suppressed the increase in mono-ADPr indicating that ADPr signal is a result of PARP14 catalytic activity. The ADPr pattern seen upon overexpression of the PARP14 MD1 mutant suggests that PARP14 modifies several different proteins in the cells while the hydrolytic activity of MD1 removes these modifications. A pull-down experiment further suggested that the signal around 200 kDa belongs to automodified PARP14. Similar results were also seen when PARP14 WT or mutants were overexpressed in U2OS cells. PARP14 mutated in MD1 showed the greatest increase in mono-ADPr with the major targets also being PARP14 itself (Fig. 4B). No induction of ADPr was observed at 50 kDa using the MD2 mutant in U2OS cells. Addition of PARG inhibitor did not affect the PARP14 signal suggesting that there is no major extension of PARP14-derived mono-ADPr into polymers (Fig. 4C). Taken together, these results suggest that PARP14 is a highly active mono-ADP-ribosyl transferase for proteins, but the level of ADPr is tightly regulated by its own hydrolytic macrodomain.

**Figure 4:**
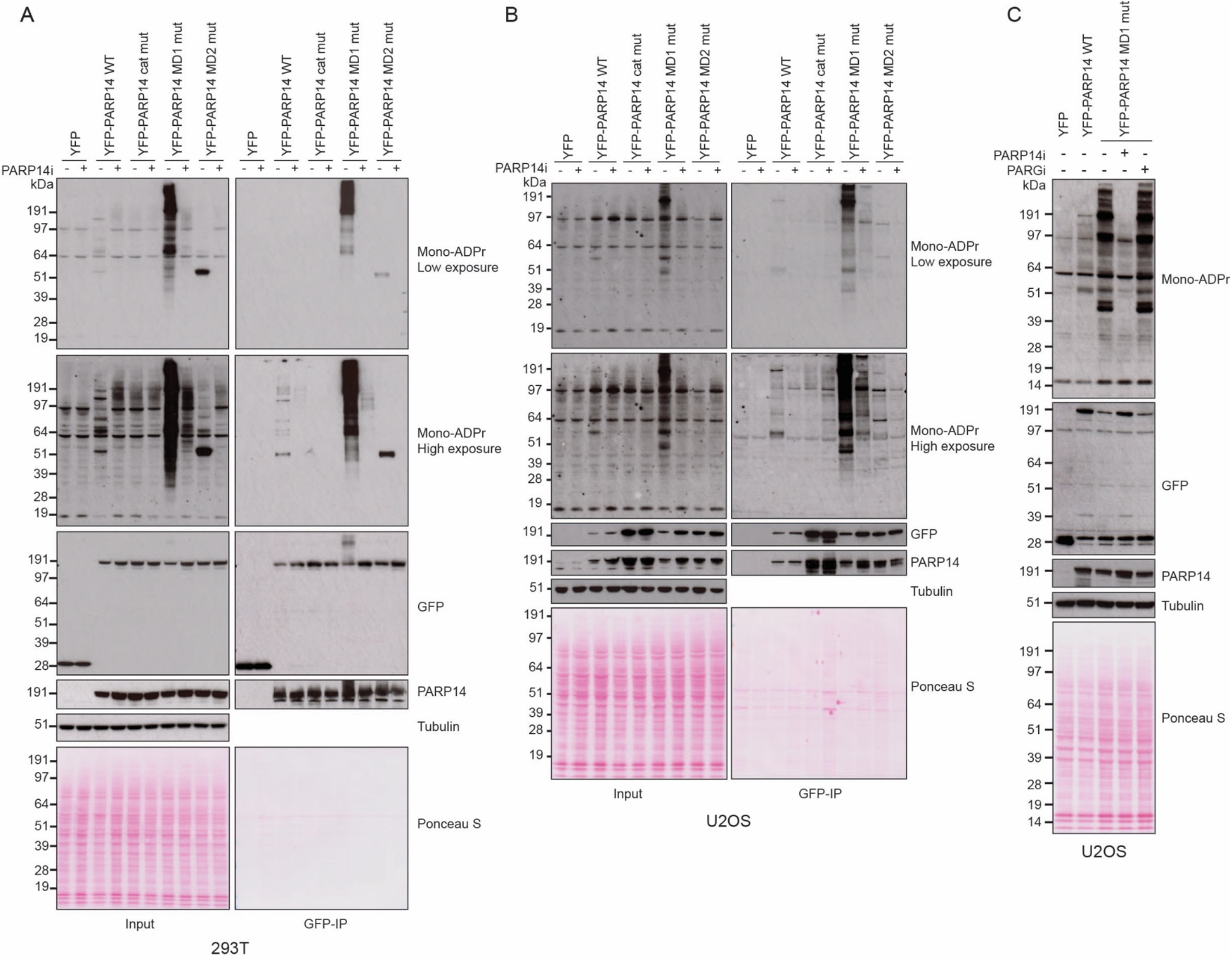
PARP14 ADPr is reversed by its own macrodomain 1. **(A)** HEK293T or **(B)** U2OS cells were transfected with the indicated plasmids in the presence or absence of PARP14 inhibitor (PARP14i). Cell lysates and GFP-immunoprecipitations (GFP-IP) were examined by western blotting using the indicated antibodies. **(C)** U2OS cells were transfected with the indicated plasmids in the presence of PARP14 or PARG inhibitor. Cell lysates were examined by western blotting with the indicated antibodies. For all blots, tubulin was used as a loading control.

### PARP14-derived ADP-ribosylation can be reversed by hydrolytic macrodomains

While PARP14 MD1 appears to be the dominant hydrolase controlling PARP14-catalysed ADPr in cells, we also tested whether PARP14 ADPr could be reversed by several other human hydrolases in a cellular context. For this, we co-expressed the PARP14 MD1 mutant, which shows the strongest increase in mono-ADPr signal upon overexpression, together with different FLAG-tagged human hydrolases (Fig. 5). Firstly, we confirmed a strong increase in mono-ADPr upon expression of PARP14 MD1 mutant, and that this could be inhibited by treatment with PARP14 inhibitor. Next, we compared PARP14-dependent ADPr in the presence of the human hydrolases MacroD1, MacroD2, TARG1 and PARG. While MacroD1 could quite efficiently remove PARP14 auto- and trans-ADP-ribosylation, the activity of MacroD2 and TARG1 could only modestly remove PARP14-dependent ADPr. Finally, PARG overexpression did not significantly reduce ADP-ribose suggesting that PARP14 catalyses largely mono-ADPr as previously suggested (*3, 38*). These results are consistent with our *in vitro* data (Fig. 2D) showing that PARP14 auto- and trans-modification targets primarily acidic residues for mono-ADPr and that its modifications are reversed by ADP-ribosylhydrolases with activity towards acidic residues such as PARP14 MD1 and MacroD1 (*45*).

**Figure 5:**
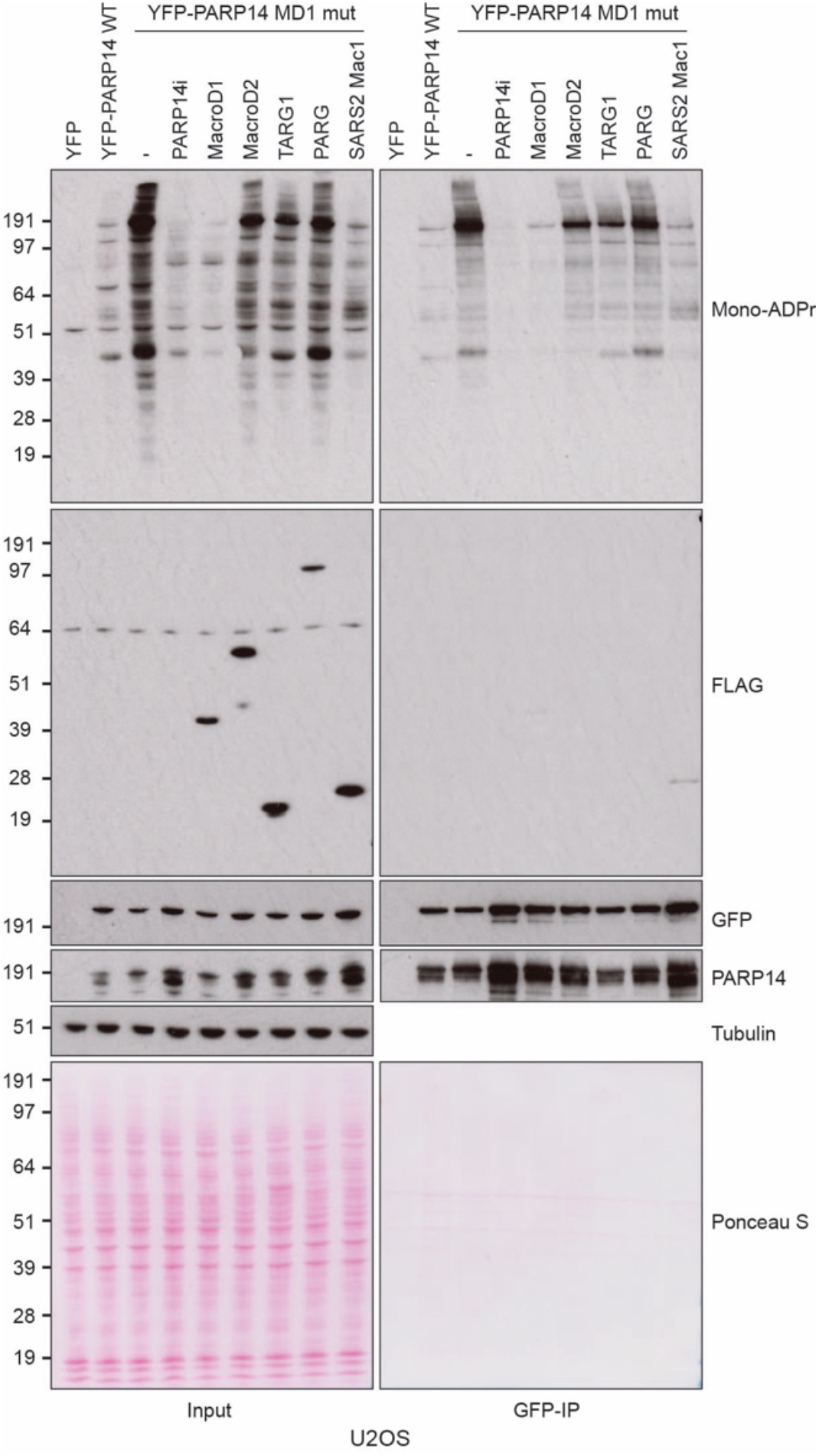
MacroD1 and SAR2-CoV Mac1 can reverse PARP14 ADPr. U2OS cells were transfected with the indicated PARP14 plasmid together in the presence or absence of PARP14i or with FLAG-tagged MacroD1, MacroD2, TARG1, PARG or SARS2 Mac1. Cell lysates and GFP-immunoprecipitations (GFP-IP) were examined by western blotting using the indicated antibodies. Tubulin was used as a loading control.

Given that SARS2 Mac1 hydrolyses PARP14 ADPr *in vitro* (Fig. 2 and (*38*)) we wanted to examine if this also held true in a cellular context. When we co-expressed SARS2 Mac1 together with PARP14 MD1 mutant, we saw a dramatic reduction in PARP14-derived ADPr (Fig. 5, lane 9), again confirming our results that these two hydrolases, PARP14 MD1 and SARS2 Mac1, can both remove ADPr catalysed by PARP14. This supports the available genetic data showing that PARP14 and coronaviral Mac1 acts as a pair where PARP14-driven ADPr acts to suppress virus proliferation while SARS2 Mac1 counteracts PARP14 antiviral activity through ADP-ribosyl hydrolysis (*26*).

### PARP13 is a target of both ADP-ribosyl transferase and hydrolase activity of PARP14

Finally, we sought to examine the hydrolytic activity of PARP14 MD1 on a known target of PARP14 ADPr. It has previously been reported that PARP14 can modify another antiviral PARP, i.e., PARP13 (*46*). To test this, we co-expressed GFP-tagged PARP13 with PARP14 WT or PARP14 MD1 mutant (Fig. 6). We observed a strong induction of ADPr of a protein the size of GFP-PARP13 in lysates of cells expressing PARP14 MD1 mutant. We then performed immunoprecipitation to pulldown GFP-PARP13 and examined its ADP ribosylation state. We found that expression of GPF-PARP13 together with WT PARP14 could modestly increase PARP13 mono-ADPr levels compared to expression with YFP alone. However, PARP13 ADP-ribosylation was greatly increased upon expression of the PARP14 MD1 mutant. Taken together, this confirms that PARP14 ADP-ribosylates PARP13 in cells, and that the hydrolytic macrodomain MD1 of PARP14 reverses this ADP-ribosylation.

**Figure 6:**
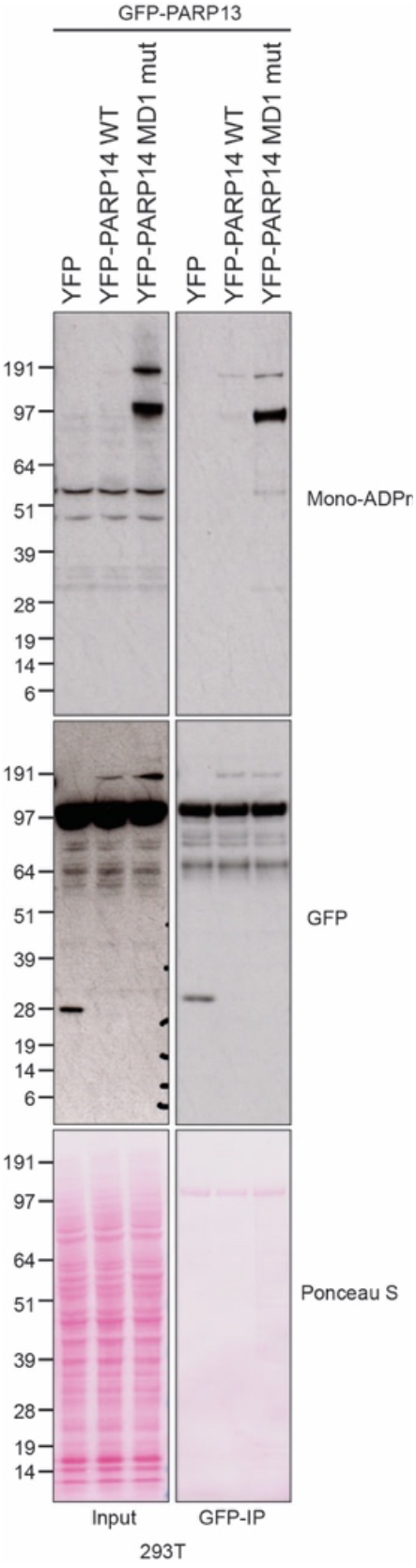
PARP14 acts to add and remove ADPr on PARP13. HEK293T cells were co-transfected with GFP-PARP13 and the indicated YFP plasmid. Cell lysates and GFP-immunoprecipitations (GFP-IP) were examined by western blotting using the indicated antibodies. Ponceau S staining was used to indicate equal loading.

## Discussion

PARP14 is involved in the regulation of several cellular processes including DNA repair, immune and antiviral response, and RNA stability (*26, 27, 29, 30, 33-35, 42, 47*). However, it is still largely unclear how PARP14 activity is achieved and regulated. While PARP14 macrodomain 2 and 3 have been identified as binders of mono-ADPr substrates (*48*), the function of the first macrodomain has not yet been characterised. Our biochemical and cellular data demonstrate that the PARP14 MD1 possesses hydrolytic activity and is the main cellular enzyme that controls the levels of PARP14 ADPr. This represents a rare example of an ADP-ribosyl transferase that also possesses ADP-ribosyl hydrolase activity. Analogously, PARP9 MD1 also possess ADP-ribosyl hydrolase activity, but PARP9 lack the transferase activity (*3*). PARP9 MD1 may contribute to the control of PARP14 ADPr levels but it could equally control the ADPr levels of other transferases, or under specific cellular conditions. However, the fact that PARP9 and PARP14 appear to be in the same complex and that they are expressed from the same genomic region (Chromosome 3q21.1) suggest that the activities of these two PARPs regulate the same pathways (*34, 42*). Altogether, our findings reveal two new ADP-ribosyl hydrolase enzymes that are present in humans and many higher organisms in addition to the already known six ADP-hydrolases -PARG, ARH1/3, MacroD1/2 and TARG1 (*11*). Our discovery that PARP14 MD1 and PARP9 MD1 is specific for glutamate ADPr may suggest some redundancy with the other Glu-ADPr targeting enzymes TARG1, MacroD1 and MacroD2 (*19, 43, 49*).

Our data shows that mutating PARP14 MD2 in cells leads to the accumulation of a single, specific ADP-ribosylated protein (Fig. 4). While we did not observe any activity of MD2 against the model ADP-ribosylated substrates (Fig. 2 and 3), there are several possibilities for the prominence of this modification. Firstly, it is possible that MD2 has catalytic activity against some specific protein(s). Alternatively, the mutation of MD2 may (allo)sterically preclude binding and hydrolysis of ADPr by MD1 on this substrate. Finally, it is possible that PARP14 is recruited to the specific protein by MD2. Thus, mutating MD2 abolishes PARP14 recruitment and subsequent hydrolysis by MD1.

PARP14 has been shown to play important roles in immunity and replication stress (*33, 35*). PARP14 MD1 and PARP9 MD1 are the most closely related human enzyme to SARS2 Nsp3 Mac1, more closely in fact than to the human paralogues MacroD1 and TARG1 (Fig. 1). It is conceivable that coronaviruses and some other viruses bearing macrodomains such as alphaviruses and hepatitis E (*37, 50*), highjacked MD1 at some point in evolution and now use it to oppose PARP14 ADPr antiviral activities. However, it cannot be ruled out that the specificities of these two macrodomains at least on some ADPr sites in macromolecules is different and may have diverged through evolution in the host–virus arms race. Linking to this, both PARP14 and other antiviral PARPs and viral macrodomains are under positive natural selection (*28, 38, 51*). Recent genetic data convincingly shows that PARP14 functions as an antiviral enzyme in a murine coronavirus model and that Mac1 counteracts this activity, antagonising the interferon response and enabling viral replication (*26, 36, 52*).

Our data also shows that PARP14 can reversibly ADP-ribosylate another antiviral PARP, PARP13 (Figure 6). Although not catalytically active, PARP13 has been implicated in inhibiting the replication of multiple classes of viruses including retroviruses (*53*), alphaviruses (*24, 54*), flaviviruses and filoviruses (*55*). It is tempting to speculate that PARP14 induced ADPr of PARP13 might be involved in the regulation of PARP13 activity, namely, increased ADPr of PARP13 might affect its stability and/or binding affinity. However, further studies are required to understand the crosstalk between these two antiviral PARPs.

PARP14 has been associated with the development of inflammatory diseases such as allergic asthma (*30*) and inflammatory arterial diseases (*42*) as well as various types of cancer including B-cell lymphoma (*56*), multiple myeloma (*57*), prostate cancer (*58*) and hepatocellular carcinoma (*59*). Therefore, PARP14 has emerged as a potential drug target prompting the development of several PARP14 inhibitors (*39, 60-62*), although none targeting PARP14 macrodomain 1. The discovery of the hydrolytic activity of PARP14, and PARP9, macrodomain 1 potentially presents a new druggable target, together with SARS2 Nsp3 Mac1, that could be used to manipulate PARP9 and PARP14 dependent pathways or function as potent antivirals.

Altogether, our data identify PARP14 as a complex protein with the specific domains that enables it to function as a writer (ART), reader (MD2, MD3) (*48*) and eraser (MD1) of ADP-ribosylation in addition to nucleic acid binding domains (Fig. 1A). This together with the interplay of the PARP9/DTX3L complex and ubiquitylation signaling (*42, 63*) is expected to have far reaching consequences on the physiology of the cell and human disease.

## Materials and Methods

### Hydrolase activity analysis using luminescence detection of ADPr

The ADP-ribosylated peptides were chemically synthesized (Table 1). Serine ADPr and Arginine-ADPr peptides were synthesized as previously described (*64, 65*). Chemical synthesis of the Glu-ADPr peptide will be described elsewhere. The hydrolytic assay against the ADP-ribosylated peptides was performed as previously described (*66*). Briefly, 10 μM substrate peptides (Arginine-ADPr, Serine-ADPr or Glutamate-ADPr) were hydrolysed using 1 μM PARP14 MD1, PARP14 MD1mut, PARP14 MD2, PARP14 MD3, PARP9 MD1, PARP9 MD2 or SARS CoV2 Mac1.

**Table 1.** Peptides used in this study

| Peptide | Sequence |
| --- | --- |
| Arginine | Ac-GR(ADPr)LIFAG-OH |
| Serine | Ac-PAKS(ADPr)APAPKKG-NH<sub>2</sub> |
| Glutamate | H-AAPVE(ADPr)VVAPR-NH<sub>2</sub> |

ARH1, ARH3 and MacroD1 served as positive controls for Arginine-ADPr, Serine-ADPr and Glutamate-ADPr respectively. Hydrolysis was carried out for 1 hour at 30 °C in assay buffer (50 mM Tris-HCl, pH 7.5, 200 mM NaCl, 10 mM MgCl_2_, 1 mM DTT and 0.2 μM NUDT5) for arginine-ADPr and serine-ADPr and (50 mM PIPES, pH 6.9, 200 mM NaCl, 10 mM MgCl_2_, 1 mM DTT and 0.2 μM NUDT5) for glutamate-ADPr. The reactions were analysed using the AMP-Glo™ assay kit (Promega) following manufacturers recommendations. Luminescence was read using a SpectraMax M5 plate reader with the SoftMax Pro software (Molecular Devices). Data were analysed using GraphPad Prism.

### *In vitro* protein (ADP-ribosyl) hydrolase assay

PARP14 WWE-CAT (1 μM) with or without PARP14 MD3 (2 μM) was incubated with 50 μM NAD^+^ (spiked with ^32^P NAD^+^) in reaction buffer (50 mM Tris-HCl, pH 8.0, 100 mM NaCl, 2 mM MgCl_2_). Reactions were incubated at 37 °C for 3 hours, then stopped by addition of 0.1 μM PARP14i. Next, ADP-ribosylated substrates were incubated with PARP14 and PARP9 macrodomains (2 μM) for 1 hour. The reactions were subsequently stopped by the addition of 4X LDS sample buffer (Life Technologies) and incubation at 95 °C for 5 min. Samples were then analysed by SDS-PAGE and autoradiography.

### *In vitro* DNA and RNA (ADP-ribosyl) hydrolase assay

DNA and RNA (ADP-ribosyl) hydrolase assays were carried out as described previously (*21*). All buffers were prepared using DNase/RNase free water and filter sterilized before use. Briefly, 0.25 μM Cy3 labelled RNA or DNA (Table 2), was mixed with 500 μM NAD^+^ and 2 μM PARP14 WWE-CAT. Reactions were incubated for 1 hour at 30 °C. The ADP-ribosylation reaction was terminated by the addition of 0.1 μM PARP14i. Hydrolysis of the ADP-ribosylated DNA or RNA was initiated by the addition of 4 μM macrodomains from PARP14, PARP9 or SARS2 Mac1 followed by incubation of the reactions for 30 mins at 30 °C. Hydrolysis was stopped by adding 50 ng/μL Proteinase K and 0.15% SDS followed by incubation for 30 min at 50 °C. 2X TBE urea sample buffer (8 M urea, 20 μM EDTA pH 8.0, 2 μM Tris-HCl pH 7.5 and bromophenol blue) was subsequently added and the samples were incubated at 95 °C for 3 mins. The samples were run on a prerun denaturing urea PAGE gel (20% (w/v) polyacrylamide, 8 M urea and 1X TBE) at 7 W/gel in 0.5X TBE. The gels were visualized with laser excitation for Cy3 at 532 nM using a PharosFX Molecular Imager (BioRad).

**Table 2.**
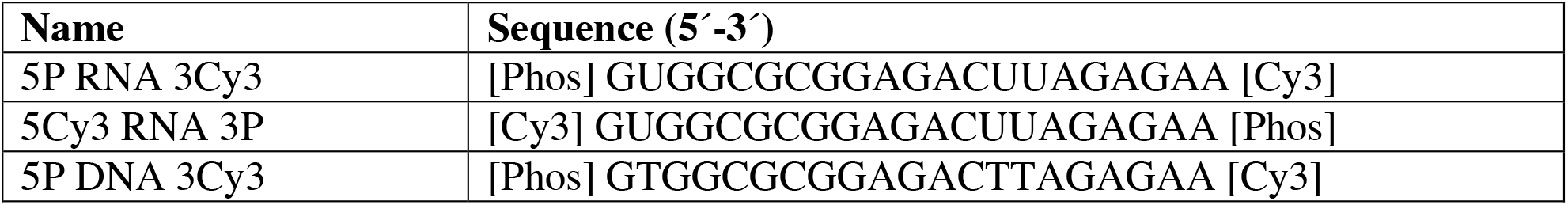
Oligos used in this study.

### Plasmids and mutagenesis

Full-length PARP14 cloning was performed by Gateway® cloning (Invitrogen) according to the manufacturer’s instructions. 300 ng of PARP14-encoding pEZ-M11 mammalian expression vector obtained from GeneCopoeia were directly set-up for BP recombination reaction with pDONR™221 vector without PARP14 insert amplification. The reaction was stopped by adding 1 μL Proteinase K (Invitrogen) and after incubation for 10 min at 37 °C. Competent Stable *E. coli* (NEB) were transformed with 2 μL of the BP reaction mix. For transfer into the destination vector, 100 ng of positive pENTR clone DNA was incubated with 100 ng of pDEST-N-YFP/FRT/TO pcDNA5 and LR Clonase™ enzyme mix for 2 hours at RT. Plasmid DNA was isolated for positive clones and verified by Sanger sequencing.

PARP14 and PARP9 macrodomains were cloned into a pNIC28-Bsa4 vector which adds an N-terminal His_6_-TEV cleavage site to the proteins to aid protein purification.

PARP14 point mutations were introduced through site-directed mutagenesis PCR using the QuickChange Lightning kit (Agilent) with primers described in Table 3 and confirmed by Sanger sequencing.

**Table 3:**
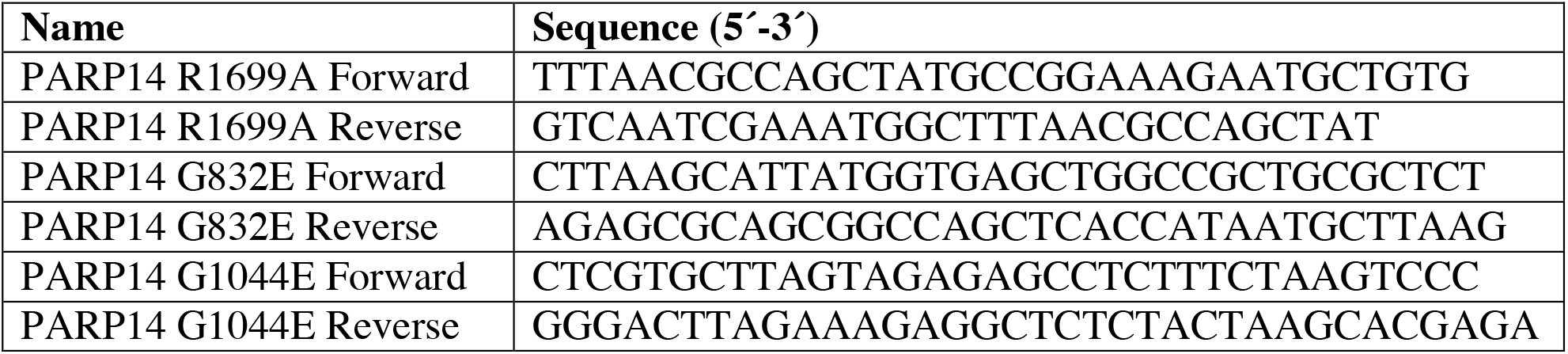
Primers used in this study

Mammalian expression vectors encoding FLAG-MacroD1, FLAG-MacroD2, FLAG-PARG, FLAG-SARS2 Mac1, and GFP-PARP13 were generated by gateway cloning as described previously (*67*).

### Protein expression and purification

BL21(DE3)-R3-pRARE cells were transformed with PARP14 and PARP9 macrodomain encoding constructs and grown at 37 °C in LB medium supplemented with appropriate antibiotics until OD_600_0.5– 0.6, then cooled to 18 °C and supplemented with 0.5 mM IPTG at an OD_600_ of 0.8 to induce protein expression overnight. Cells were harvested by centrifugation, resuspended in lysis buffer (50 mM HEPES pH 7.5, 500 mM NaCl, 20 mM imidazole, 5 % glycerol, 0.5 mM tris(2-carboxyethyl)phosphine [TCEP], 1:2,000 Calbiochem protease inhibitor cocktail set III) and lysed by sonication. Proteins were purified by Ni^2+^-NTA chromatography (Jena Bioscience) and eluted stepwise in binding buffer with 40–250 mM imidazole. Proteins were further purified by size-exclusion chromatography (SEC) (Superdex 75, GE Healthcare) in a buffer consisting of 50 mM HEPES pH 7.5, 300 mM NaCl, 5 % glycerol, 0.5 mM TCEP. PARP14 MD1 was additionally purified by ion exchange chromatography using a HiTrap™ 5 mL SP HP (GE Healthcare) equilibrated in 25 mM HEPES pH 7.5, 75 mM NaCl, 0.5 mM TCEP. The purity of protein preparations were assessed using SDS-PAGE and CBB staining and aliquots were stored at -80 °C until use.

### Cell culture

Human U2OS osteosarcoma (ATCC HTB-96) and embryonic kidney 293T (ATCC CRL-3216) cell lines were purchased from American Type Culture Collection (ATCC). Cells were grown in Dulbecco’s modified Eagle’s medium (DMEM) (Sigma) supplemented with 10% fetal bovine serum (FBS) (GIBCO) and penicillin-streptomycin (100 U/mL, GIBCO). All cell lines were cultured in a humidified atmosphere at 37°C with 5% CO_2_. HEK293T and U2OS cells were plated in 10-cm dishes 24 h before cells were transfected with the indicated plasmids. HEK293T cells were transfected using Polyfect (QIAGEN), while U2OS cells were transfected using TransIT-LT1 Transfection Reagent (Mirus Bio), according to the manufacturer’s protocol. Cells were treated with DMSO, 0.5 μM PARP14i (RBN012759, MedChemExpress) or 5 μM PARGi (PDD00017273, Sigma) for 24 h.

### Immunoprecipitation

HEK293T and U2OS cells were collected 24 or 48 h post-transfection and washed two times with PBS. Cells were lysed with Triton X-100 lysis buffer (50 mM Tris-HCl pH 8.0, 100 mM NaCl, 1% Triton X-100) supplemented with 5 mM MgCl_2_, 0.1% Benzonase (Sigma), protease and phosphatase inhibitors (Roche), Olaparib (Cayman Chemical; 1 μM for U2OS; 2 μM for 293T cells), 1 μM PARGi PDD00017273 (Sigma) for 30 min at 4°C. Protein concentrations were determined by Bradford Protein Assay (Bio-Rad) and normalised for equal protein amounts. Cell lysates were incubated with GFP-Trap magnetic agarose beads (ChromTek) on an orbital rotator for 2 h at 4°C. Beads were pelleted using a magnetic separation rack and washed five times with Triton X-100 lysis buffer (50 mM Tris-HCl pH 8.0, 800 mM NaCl, 1% Triton X-100). Proteins were eluted with 2x NuPAGE LDS sample buffer (Invitrogen) supplemented with DTT (Sigma-Aldrich), boiled for 5 minutes at 95°C and analysed by western blotting.

### Western blotting

Cells were lysed and protein concentration was measured as described above. Proteins were boiled in 1x NuPAGE LDS sample buffer (Invitrogen) with 60 mM DTT (Sigma-Aldrich) and resolved on NuPAGE Novex 4%–12% Bis-Tris gels (Invitrogen) in 1x NuPAGE MOPS SDS Running Buffer (Invitrogen) at 150 V. Proteins were transferred onto nitrocellulose membranes (Bio-Rad) using Trans-Blot Turbo Transfer System (Bio-Rad). The membranes were stained with Ponceau S Staining Solution (Thermo Fisher) to check the transfer quality, rinsed with water, and blocked in 5% (w/v) non-fat dried milk in PBS buffer with 0.1% (v/v) Tween 20 (PBST) for 1 hour at room temperature. This was followed by overnight incubation with primary antibody as indicated in Table 4 at 4°C. The next day, membranes were washed in PBST and incubated with HRP-conjugated antibodies at room temperature for 1 h. Membranes were visualised on Hyperfilm ECL films (Cytiva) after adding Pierce ECL Western Blotting Substrate (Thermo Fisher).

**Table 4:** Antibodies used in this study

| Target | Host | Company | Reference | Dilution in WB |
| --- | --- | --- | --- | --- |
| GFP | Rabbit | Abcam | Ab290 | 1:3000 |
| PARP14 | Rabbit | Abcam | Ab229756 | 1:1000 |
| β-tubulin | Rabbit | Abcam | Ab6046 | 1:2000 |
| FLAG | Rabbit | Sigma-Aldrich | F7425 | 1:2000 |
| HRP conjugated anti-Mono-ADPr | - | - | Gift from Ivan Matic | 1:1000 |
| HRP-conjugated anti-mouse | Goat | Agilent | P0447 | 1:3000 |
| HRP-conjugated anti-rabbit | Swine | Agilent | P0399 | 1:3000 |

### Sequence and phylogenetic analysis

For multiple-sequence alignments, JalView v2 (*68*) and MAFFT7 (*69*) was used. The phylogenetic tree of macrodomains was generated with SplitsTree4 (v4.15.1) (*70*) using the Neighbour-Joining (NJ) method and confidence levels estimated using 1000 cycles of the bootstrap method. Pairwise identities were determined using the Needleman–Wunsch algorithm implemented as part of the EMBL-EBI search and sequence analysis server (*71*). Structural alignments and analyses, as well as figure preparation, were carried out using PyMol (Molecular Graphics System, Version 2.3.3 Schrödinger, LLC). PARP domain architecture was visualised using IBS illustrator (*72*).

## Acknowledgements

We would like to thank Ivan Matic for his kind gift of HRP-conjugated anti-mono-ADPr antibody. We would like to thank Andreja Mikoc for critical reading of our manuscript.

## Funding

Biotechnology and Biological Sciences Research Council (BB/R007195/1 and BB/W016613/1) (IA)

Wellcome Trust (210634) (IA)

Oxford University Challenge Seed Fund (USCF 456) (IA)

Edward Penley Abraham Research Fund (IA)

Ovarian Cancer Research Alliance (813369) (IA)

Research Council of Norway (315849) (ØS)

Wellcome Trust (223107) (IA and SS)

Swedish Research Council (2019-04871) (HS)

MRC CDA (MR/X007472/1) (JGMR)

## Author contributions

Conceptualization: IA Methodology: DVF, SW, HS

Investigation: NĐ, ØS, DM, KZ, MS, CC, PK, JGMR, DB, SW, RS

Visualization: NĐ, ØS, KZ, MS, RS

Supervision: SS, DVF, DA, IA

Writing—original draft: RS, NĐ, ØS, IA

Writing—review & editing: NĐ, ØS, DM, KZ, MS, PK, JGMR, DB, HS, DVF, SS, RS, DA, IA

